# Framework to simulate gene regulatory networks with stochastic molecular kinetics and to infer steady-state network structure

**DOI:** 10.1101/872374

**Authors:** Johannes Hettich, J. Christof M. Gebhardt

## Abstract

**Background:** The temporal progression of many fundamental processes in cells and organisms, including homeostasis, differentiation and development, are governed by gene regulatory networks (GRNs). GRNs balance fluctuations in the output of their genes, which trace back to the stochasticity of molecular interactions. Although highly desirable to understand life processes, predicting the temporal progression of gene products within a GRN is challenging when considering stochastic events such as transcription factor – DNA interactions or protein production and degradation.

**Results:** We report CaiNet, a fast computer-aided interactive network simulation environment optimized to set up, simulate and infer GRNs at molecular detail. In our approach, we consider each network element to be isolated from other elements during small time intervals, after which we synchronize molecule numbers between all network elements. Thereby, the temporal behaviour of network elements is decoupled and can be treated by local stochastic or deterministic solutions. We demonstrate the working principle of the modular approach of CaiNet with a repressive gene cascade comprising four genes. By considering a deterministic time evolution within each time interval for all elements, our method approaches the solution of the system of deterministic differential equations associated with the GRN. By allowing genes to stochastically switch between on and off states or by considering stochastic production of gene outputs, we are able to include increasing levels of stochastic detail and approximate the solution of a Gillespie simulation. Notably, our modular approach further allows for a simple consideration of deterministic delays. We further infer relevant regulatory connections and steady-state parameters of a GRN of up to ten genes from steady-state measurements by identifying each gene of the network with a single perceptron in an artificial neuronal network and using a gradient decent method originally designed to train recurrent neural networks.

**Conclusion:** CaiNet constitutes a user-friendly framework to simulate GRNs at molecular detail and to infer the topology and steady-state parameters of GRNs. Thus, it should prove helpful to analyze or predict the temporal progression of reaction networks or GRNs in cellular and organismic biology. CaiNet is freely available at https://gitlab.com/GebhardtLab/CaiNet.

## Background

Dynamics and progression of many fundamental processes in cells and organisms, including metabolism^1,2^, the cell cycle^3^, the circadian clock^4^, differentiation^5,6^ and development^7^ are governed by GRNs. These networks generally control the activity of genes by regulatory motifs such as feedback or feed forward loops^8,9^, which ensure spatially and temporally controlled gene expression. A striking property of gene transcription is its distinct stochastic behavior. Rather than producing their product continuously, most genes stochastically transit from an inactive state into an active state where they produce the gene product^10,11^. During the on-time, which often is short compared to the off-time, a burst of expression occurs that might significantly increases the expression level. In combination with stochastic production (birth) and degradation (death) processes, this leads to complex stochastic trajectories of gene products. This noise of gene expression was shown to play an important role in decision making within GRNs^12–15^. Furthermore, theoretical work suggests that this stochastic behavior can influence the functionality of networks such as the circadian clock^16,17^ and the pluripotency network^18,19^.

To model and simulate the average temporal behavior of GRNs, ordinary differential equations (ODEs) and corresponding solvers are well established^20,21^. To add stochasticity, the Langevin method adds normal distributed gene product noise to the system, assuming that all reactions in the system went through multiple discrete reaction events during a single simulation time step^22^. While this method achieves a reasonable description of systems with large molecule numbers, it does not explicitly account for gene on/off switching. An exact approach to simulate stochastic processes is the Gillespie direct method^23^. In this method, two random numbers drawn from a probability distribution comprising all possible reactions in the network determine the time point and the type of the next reaction event. In this approach, the duration of a time-step is limited by the fastest occurring reaction in the network. Thus, for networks including large molecule numbers or reactions with fast rates that occur on a timescale much shorter than the total simulation time, the Gillespie direct method is computationally expensive. To mitigate this problem, the tau-leaping method has been developed^24^. There, a simulation time step may span over several reaction events. The number of reaction events that occurred during a time step is approximated by an average number. This method requires additional modifications to prevent negative particle numbers ^25^ and to calculate step sizes^26^. To further speed up simulations, hybrid approaches that combine all of the aforementioned approaches have been developed ^27–31^. These approaches treat specific reactions in the network with different algorithms. To do so, complex considerations to decouple a generic network have to be performed ^27^. Considering special cases of networks, i.e. networks with genes stochastically switching between on and off states, led to more effective dedicated hybrid stochastic deterministic simulation approaches^32^. However, a dedicated hybrid approach capable of modelling gene product synthesis in detail and providing biochemical reactions to simulate signaling pathways is still missing.

The inverse problem to simulating expression levels of GRNs with given parameters is to infer parameters and connections between genes from a set of given gene product levels, knockout data or time trajectories. A prominent approach to infer the topology of networks is based on Boolean interactions^33,34^. Another approach looks for correlation coefficients^35^ between steady state networks. Furthermore, linearized differential equations have been used to fit experimental data^36^. Using knockout data, a confidence matrix for network interactions was obtained^37^. Recent progress to the inference problem has been made using time trajectories of gene product levels^38–41^. Approaches such as WASABI^42^ infer how perturbations of gene product levels propagate through a network. However, many inference approaches indentify the relative importance of network parameters or connections, instead of yielding parameters that can directly be entered into physiological simulations.

Since genes may combine the actions of several transcription factors for their specific output behavior, they have been identified with perceptrons or neurons ^43–46^. This enabled optimizing sophisticated algorithms originally designed for artificial neural networks, such as gradient decent methods, to the inference problem ^47–49^. Using such an approach, a neural network was trained to reproduce a time-series of gene product levels^49^. During training, the neural network had to learn both how to simulate the ODE system corresponding to the gene network as well as the network topology. Thus, it remained a challenge to disentangle network parameters and topology from the coefficients of the neural network.

Here, we report a computer aided interactive gene network simulation tool (CaiNet), dedicated for simulation of GRNs including molecular kinetics and for inference of steady-state network parameters. We simplified the simulations by determining the temporal evolution of each network element isolated from all other network elements between fixed time steps, after which the gene product levels are synchronized within the GRN. This modular approach enabled us to include local analytical solutions of the temporal evolution of a network element such as a gene, thereby speeding up simulation time. For genes, we included local analytical solutions to the time behavior for various different regulatory promoter motifs and variable numbers of rate-limiting steps in gene product synthesis. In addition, we provided local analytical solutions for elementary biochemical reactions such as dimerization, that can be combined to model complex biochemical reaction networks. We tested CaiNet by comparing its simulation of a repressive gene cascade of four genes to the solution of the global, network-wide ODEs by an ODE solver and to a full stochastic Gillespie simulation. For parameter inference, CaiNet uses a recurrent network approach to directly infer steady-state parameters. We tested CaiNet for up to 10 densely connected genes and find that CaiNet is able to recover the network topology and the network parameters well. The combination of a simulation algorithm and an inference algorithm directly yielding physiological simulation parameters will remove hurdles in understanding experimental results and investigating associated GRNs.

## Results

### Setup of GRNs with the graphical user interface of CaiNet

GRNs commonly comprise several elements: external signaling inputs, biochemical reactions and regulatory motifs including several genes (Figure 1a, upper panel). In CaiNet, we characterized each element by certain structural and kinetic parameters (Figure 1b). Inputs may be any kind of molecule. Each molecule input can be assigned a time course of molecule abundance, which might be for example sinusoidal or rectangular (Figure 1c). Complex biochemical reactions can be set up by combining a certain set of elementary reactions^23^. In CaiNet, this set comprises homodimerisation, heterodimerisation and a transformation of a species by an enzyme. For example, formation of a homotetramer can be implemented by combining the homodimerization of a monomer and a subsequent homodimerization of the homodimer. We modeled genes by two states, an on-state and an off-state^50^, and a promoter with a certain number of binding sites for transcription activators or repressors (Figure 1b and 1d). The gene switches to the on-state with the effective rate *λ_eff_* upon association of activating transcription factors according to the regulatory logic of the promoter, and switches to the off-state with the effective rate *μ_eff_* upon their release (Figure 1d and Methods). Transcription repressors keep the gene in the off-state (Methods). In the on-state, the gene product is produced with the production rate *ν*. The gene product can either be understood as mRNA or as protein. In the latter case, we initially simplified the process of protein production by combining transcription and translation of mRNA into a common rate-limiting step with one production rate. However, we introduced the possibility for an additional delay to account for production processes such as splicing and translation (see below). Moreover, gene products are associated with a degradation rate.

**Figure 1:**
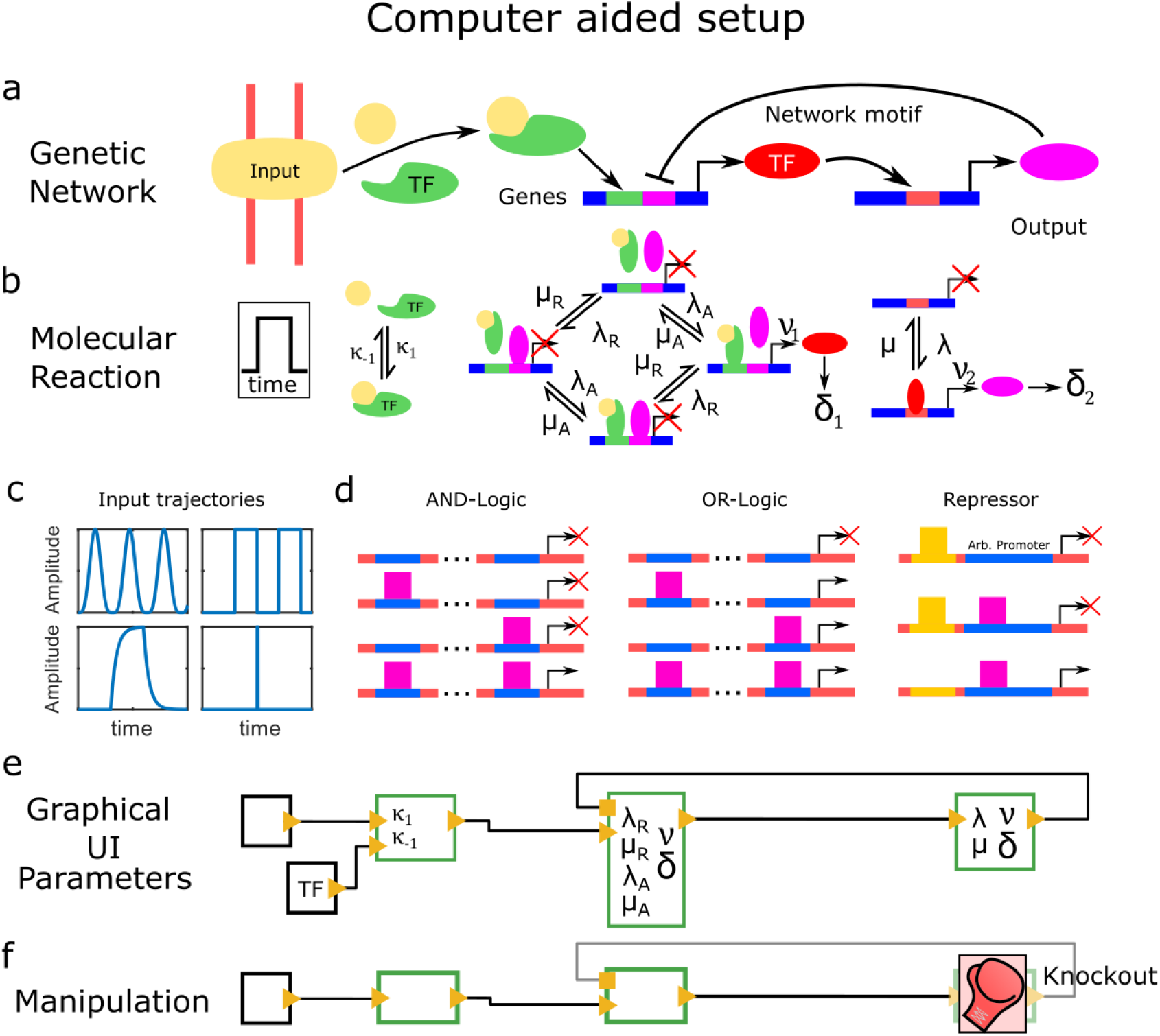
Implementation of GRNs in CaiNet. (a) Sketch of an exemplary GRN. GRNs in CaiNet may comprise elements of (extra)cellular input signals, elements of basic bi-molecular reactions combined to more complex biochemical reactions and gene elements combined to regulatory motives. (b) Sketch of the reaction rates entering the exemplary GRN. CaiNet provides analytical solutions to the evolution of molecule numbers in basic biochemical reactions and analytical effective two-state solutions of molecule numbers of different promoter structures of genes, which all are included into the unique modular simulation approach of CaiNet. The user can assign time profiles of input elements and reaction rates of bi-molecular reactions and genes. (c) Exemplary time trajectories for input elements. (d) Sketch of promoter structures implemented in CaiNet. (e) Sketch of the layout of elements including their kinetic parameters of the exemplary GRN in (b) set up in CaiNet via a GUI. (f) Sketch of the knockout tool applied to the exemplary GRN. The parameters of elements can be altered to simulate experimental conditions such as knockdown or knockout experiments.

We designed a graphical user interface (GUI) for CaiNet to facilitate setting up complex GRNs (Figure 1e), inspired by an effort to find general and understandable representations of biological networks^51^. With this GUI, icons representing the network elements ‘input’, ‘bimolecular reaction’ and ‘gene’ can be connected intuitively using activating or inhibiting links represented by wires. For each network element, relevant structural and kinetic parameters can be defined (Figure 1e and Table 1). In addition, we implemented means to manipulate a genetic network by knocking down one or more genes (Figure 1f). For a knocked down gene, the transcription rate is adjusted to a low value or zero.

**Table 1:**
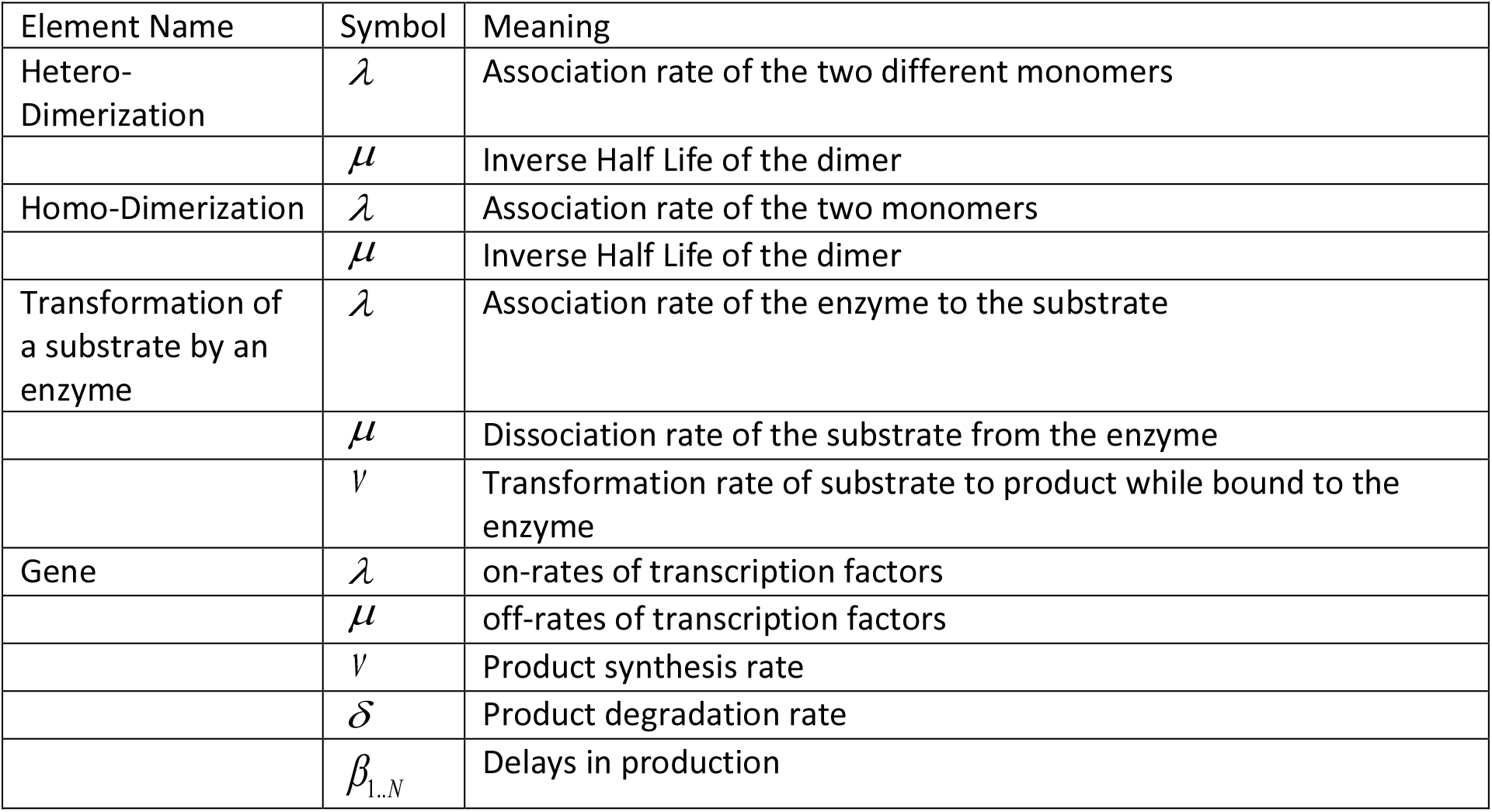
Kinetic parameters of network elements

### Computer-aided implementation of network simulations

To simulate GRNs with CaiNet, we developed a modular approach of solving the system of differential equations associated with the GRN or of simulating the stochastic behaviour of genes and of the numbers of gene products. At the heart of this modularisation lies the assumption that changes in molecule numbers within the GRN are small during a time step *Δt* and that it is sufficient to synchronize the numbers of molecules between all elements of the network only after each time step *Δt*. According to this assumption, each network element can be treated as if it was isolated from other network elements during *Δt*. Thus, for each *Δt*, CaiNet iterates through all network elements and calculates the local changes in molecule abundance over time for each element using the molecule numbers communicated to the element at the previous synchronization step and element-specific functions (Methods, pseudo code). We derived various such functions for bi-molecular reactions and different promoter structures of genes (Methods). After each *Δt*, CaiNet collects the updated molecule numbers from each network element and synchronizes the new network-wide numbers between all elements. Importantly, our modular enabled simulating different scenarios with increasing level of stochastic detail, ranging from an approximate solution of the system of deterministic differential equations associated with the GRN to an approximate solution of the corresponding stochastic chemical master equations.

To demonstrate the working principle of our modular approach, we chose a cascade of negative regulation comprising four genes (Figure 2a). In the cascade, each gene encodes a repressive transcription factor, which represses the subsequent gene. Such a cascade has been shown to exhibit pathologic behaviour in presence of stochasticity ^52,53^. We set the number of the first repressive transcription factor, which acts as initial input, to a constant value (high). We then determined the numbers of subsequent transcription factors during the simulation. In the following, we discuss the different scenarios considering increasing levels of stochastic detail.

**Figure 2:**
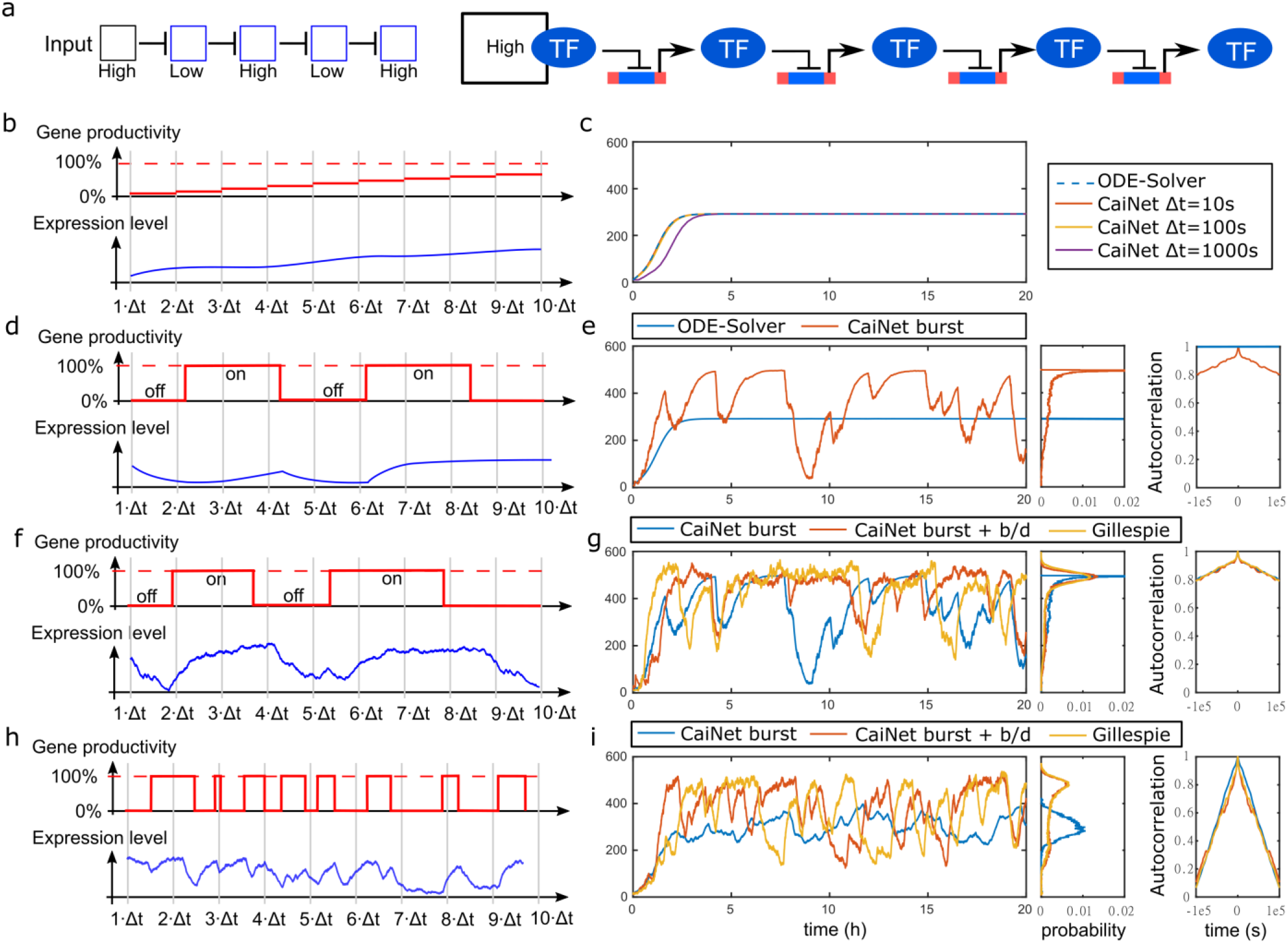
Demonstration and test of CaiNet using a repressive gene cascade. (a) Regulatory logic of a repressive gene cascade of four genes (left panel) and sketch of the corresponding GRN including transcription repressors and gene elements (right panel). The input species and each gene product repress the subsequent gene element in the GRN. (b) Sketches of the probability of activated expression of a gene (upper panel) and the corresponding gene product level (lower panel) of scenario 1 (all elements of the GRN treated deterministically) of the CaiNet simulations. The activation of a gene is constant within one synchronization time step. After each time-step, all gene product levels are synchronized and the activation probabilities of all gene elements are updated. (c) Comparison of CaiNet simulations of the repressive gene cascade according to scenario 1 performed with different synchronization time steps (red, yellow and purple lines) with the numerical solution of an ODE solver (dashed blue line). (d) Sketches of the production state of a gene (upper panel, either on or off)) and the corresponding gene product level (lower panel) of scenario 2 (gene on/off switching treated stochastically) of the CaiNet simulations. After each synchronization time-step, all gene product levels are synchronized and the effective on/off rates of all gene elements are updated. (e) *Left panel:* Comparison of a CaiNet simulation of the repressive gene cascade according to scenario 2 (red line) with the numerical solution of an ODE solver (blue line). (f) Sketches of the production state of a gene (upper panel, either on or off)) and the corresponding gene product level (lower panel) of scenario 3 (gene on/off switching and birth/death processes treated stochastically) of the CaiNet simulations. After each synchronization time-step, all gene product levels are synchronized and the effective on/off rates of all gene elements are updated. (g) *Left panel:* Comparison of a CaiNet simulation of the repressive gene cascade according to scenario 2 (blue line) and scenario 3 (red line) with a Gillespie simulation (yellow line). (h, i) As in (f, g), but with faster gene on/off switching rates. *Middle and right panels of (c, e, g, i):* Histograms and autocorrelation curves of respective gene product levels. In panels (c,e,g,i) the expression level of the last transcription factor in the cascade is shown.

Scenario 1 (Figure 2b and c): In this scenario, we implemented the approximate solution of CaiNet to the system of deterministic differential equations associated with the GRN of the repressive gene cascade (Methods, Equation(59)). We modelled the protein output of each gene by

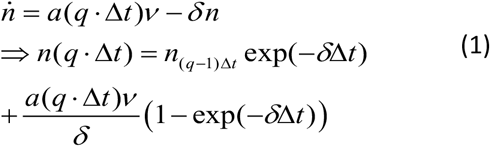

where *n* is the number of proteins, *ν* is the production rate and *δ* is the degradation rate. Gene elements directly use an analytical solution to calculate the expression level. We calculated the probability function of activated expression, *a(q Δt)*, using the expression levels from the last synchronization time point, *q Δt*, and the equilibrium constants of transcription factors of the appropriate promoter structure (Methods, Equation (13) or (16)). According to our central assumption, the probability function *a* is constant during the time interval *Δt* (Figure 2b). Thus, we could find an individual local analytical solution for each gene element in the network. Using CaiNet, we simulated the gene cascade with a fixed set of kinetic rates for the transcription factors (Table 2) and for three different time steps *Δt* (*Δt* = 10s, 100s and 1000s) (Figure 2c). If *Δt* was too large, the synchronization of transcription factor abundance and thus the change in gene activation was delayed. Hence, CaiNet’s solution for the transient behavior did not match the global, network-wide solution of the ODE solver (Methods, Equation (59)) anymore. However, the steady state level was still reproduced correctly. If *Δt* was sufficiently small, the approximation of CaiNet approached the global, network-wide solution of the system of ODEs associated with the GRN..

**Table 2:** Kinetic parameters of the repressive gene cascade

| Paramter | Scenario: Slow Switching | Scenario: Fast Switching |
| --- | --- | --- |
| **Gene 1** |  |  |
| $\lambda_A$ | 0.05 | 1 |
| $\mu_A$ | 0.05 | 1 |
| $\lambda_R$ | 4.4 | 87 |
| $\mu_R$ | 0.05 | 1 |
| $\nu$ | 0.1 | 0.1 |
| $\delta$ | 0.001 | 0.001 |
| **Gene 2** |  |  |
| $\lambda_A$ | 0.05 | 1 |
| $\mu_A$ | 0.05 | 1 |
| $\lambda_R$ | 4.1 | 82 |
| $\mu_R$ | 0.05 | 1 |
| $\nu$ | 0.1 | 0.1 |
| $\delta$ | 0.001 | 0.001 |
| **Gene 3** |  |  |
| $\lambda_A$ | 0.05 | 1 |
| $\mu_A$ | 0.05 | 1 |
| $\lambda_R$ | 12.5 | 250 |
| $\mu_R$ | 0.05 | 1 |
| $\nu$ | 0.1 | 0.1 |
| $\delta$ | 0.001 | 0.001 |
| **Gene 4** |  |  |
| $\lambda_A$ | 0.05 | 1 |
| $\mu_A$ | 0.05 | 1 |
| $\lambda_R$ | 8 | 160 |
| $\mu_R$ | 0.05 | 1 |
| $V$ | 0.5 | 0.5 |
| $\delta$ | 1e-3 | 1e-3 |

Scenario 2 (Figure 2d and e): We included that genes can stochastically switch between on and off states (Figure 2d). For simplicity, we assumed that genes were switched on with rate *λ(n_TF_*) upon binding of a transcription factor, and switched off with rate *μ* upon unbinding of the transcription factor. Thus, the switching events are Poisson processes. For every time step *Δt*, we calculated the effective on/off-rates, *λ*,*μ*, of the gene using the appropriate promoter structures (Methods Equation (14) or (17)). According to our central assumption, *λ*,*μ*, were constant within a time step *Δt*. In contrast, the gene was able to switch states within a time step. To obtain the time points of switching, *t_i_*, we drew the uniformly distributed random number *X* and used the current switching rate constant

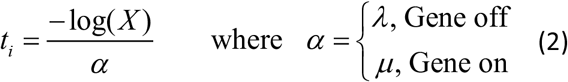

Dependent on the state of the gene, we distinguished between the two equations

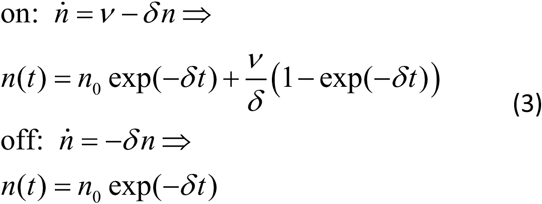

Gene elements directly used the corresponding analytical solutions to calculate the expression level. Using CaiNet, we simulated the gene cascade with a fixed set of kinetic rates for the transcription factors (Table 2) and allowed for gene switching (Figure 2e). We observed that protein abundance increased during the gene on-state and decreased while the gene was off. In combination with stochastic switching between these two states, the variance of the expression-level was significantly increased compared to the global, network-wide solution of the ODE-solver (Figure 2e). Correspondingly, while the autocorrelation of the protein output of the gene cascade was constant for the solution of the ODE-solver, the autocorrelation for the solution including switching of genes exhibited a decay since protein levels varied over time (Figure 2e).

Scenario 3 (Figure 2f and g): In addition to the switching of genes, we included discretized and stochastic production and degradation events (birth and death events) of proteins (Figure 2f). We initially determined the time points of gene switching according to Equation (2) using constant protein numbers for a given *Δt*. For each on-period of the gene with duration *τ_on_*, we determined the number of birth events *n_B_* of gene products by drawing a random number from the probability distribution corresponding to protein production:

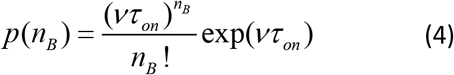

while the number of degraded gene product is determined deterministically according to Equation 3. In a period without production, the probability to find *k* proteins after the time interval *τ_off_* is given by

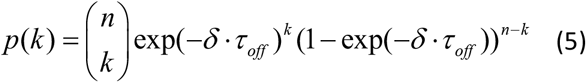

assuming there were *n* proteins at the beginning of the time interval. Using CaiNet, we simulated the gene cascade with a fixed set of kinetic rates for the transcription factors (Table 2), allowed for gene switching, and accounted for birth/death events (Figure 2g). Compared to the case including gene switching alone (scenario 2), the variance further increased and approached the variance of a Gillespie simulation of the gene cascade considering the promoter structure explicitly and including stochastic birth and death events of proteins (Methods).

We further tested the behavior of the gene cascade in a situation where the rates of gene switching were much faster than those for production (birth) and degradation (death) events (Figure 2h and i and Table 2). For scenario 2 without birth/death events of proteins, fast rates of gene switching underestimated the variance in protein levels compared to a global, network-wide Gillespie simulation including both gene switching and birth/death events (Figure 2i). This indicates that the main contribution to variance in protein levels in this situation is due to birth/death events. Accordingly, when we included both gene switching and birth/death events in CaiNet (scenario 3), the variance in protein levels well approached the variance of the global, network-wide Gillespie simulation, despite the simplified treatment of birth and death events in CaiNet. Similarly, the autocorrelation of protein output, which characterizes the temporal variation of protein levels, was similar for the protein levels obtained by CaiNet with scenario 3 and Gillespie.

To ensure sound consideration of gene product numbers in all three scenarios, we implemented that CaiNet monitors the synchronization time step and returns a warning if the change of a gene product, *Δn*, during one interval violates the criterion

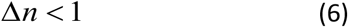

For the transcription rate *ν*, for example, this leads to the condition

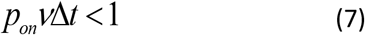

Thus, if *ν* was the fastest rate constant in the system, choosing the synchronization time step *Δt* < *ν*^−1^ ensures small changes in gene elements.

### Consideration of delays

The processes in a cell may give rise to temporal delays. For example, the process of elongation may be considered as a constant delay in gene transcription. We implemented the possibility to account for deterministic, constant delays in the response of a network element in CaiNet as a shift of the element’s output by a fixed number of synchronization steps *Δt* (Figure 3a). For example, if the output of a network element is delayed by 10 *Δt*, the element internally stores the value from ten synchronization steps and reports the delayed value during synchronization. A limitation of this approach is that delays can only be given as multiples of the synchronization time step *Δt*.

**Figure 3:**
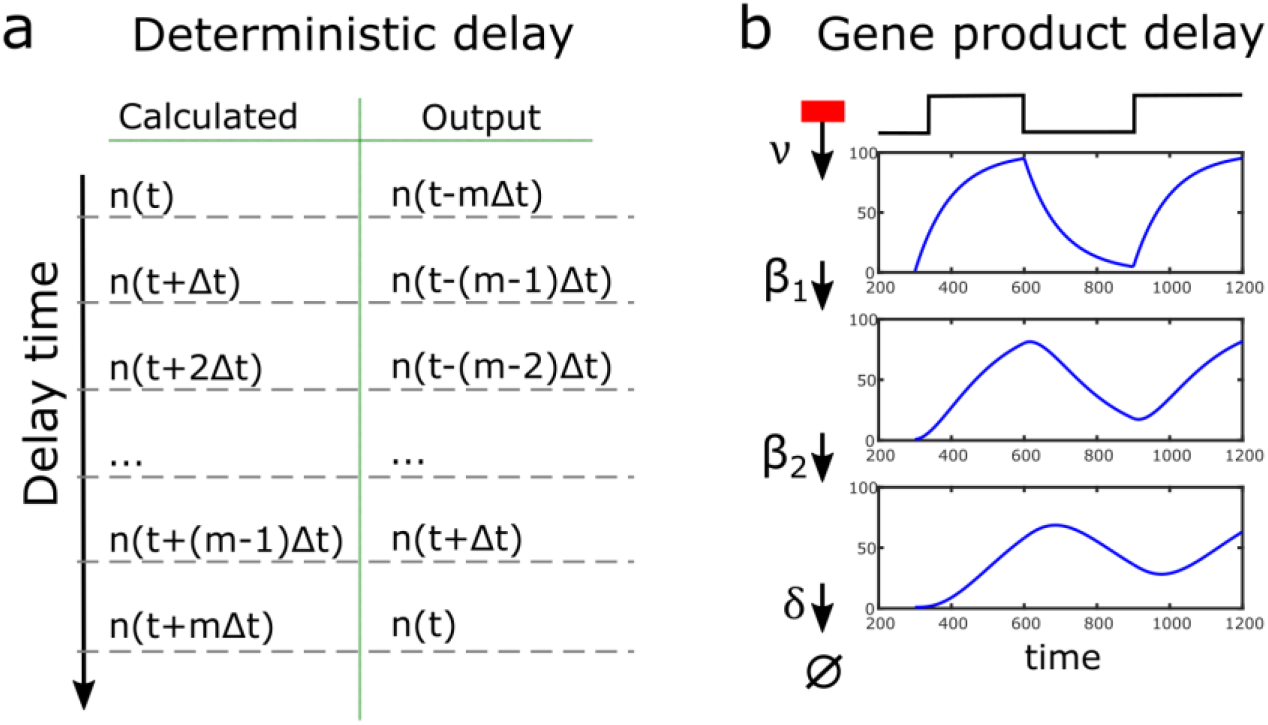
Implementation of delays in gene product synthesis in CaiNet. (a) A deterministic delay, i.e. a shift by a constant time-period, is realized by a queue. At time *t*, network elements report the expression levels from time point *t* – *m*Δ*t*, where *m*Δ*t* is the time by which the gene product synthesis is delayed. (b) Rate-limiting steps in gene product synthesis are modelled by analytical solutions of the corresponding system of ODEs. Such delays modify the shape of the time trajectory of the expression level and slightly delay the maxima of expression.

Moreover, we integrated the possibility to account for several rate-limiting steps in gene product synthesis. Such delays may occur due to transcription termination, mRNA splicing or translation. We modeled each rate-limiting step as a Poisson process with rate *β_i_*, i.e. with exponentially distributed waiting times. Thus, changes in gene activity are blurred if average waiting times are on the order of or slower than gene on/off switching processes (Figure 3b). We only simulated rate-limiting steps deterministically. Therefore, Equation (1) was replaced by an analogous equation including the appropriate number of rate-limiting steps (Methods, Equations (24) to (29)). At the synchronization time step *Δt*, the gene element returns the number of gene products of the corresponding solution.

### Consideration of biochemical reactions

We further implemented the possibility to include and combine network elements of bi-molecular reactions in CaiNet (Figure 4). To demonstrate the working principle, we chose an example of two enzymes that mutually take each others product and catalyze it to each others substrate (Figure 4a). The differential equations for the substrate *N* and the enzyme *F* that transforms the substrate *N* into the product *M* are

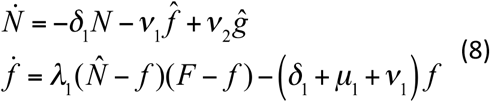

where *δ* is the degradation rate and *ν*_1_ and *ν*_2_ are the catalytic rates of the enzymes *F* and *G*. The association and dissociation rates of the substrate to the enzymes are denoted by *λ*_1_ and *μ*_1_ for *N* and *F* and by *λ*_2_ and *μ*_2_ for *M* and *G*. The differential equations for the enzyme *G* that transforms *M* back into *N* are

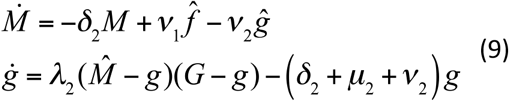

**Figure 4:**
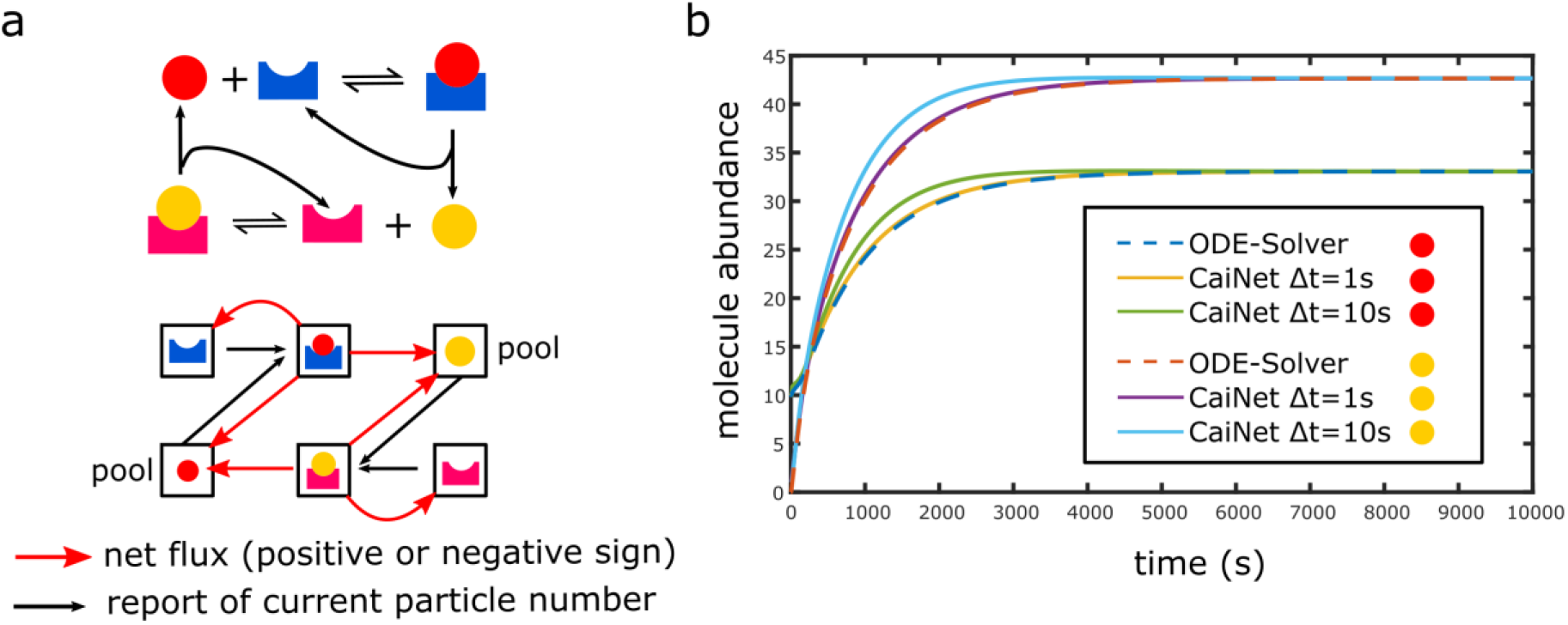
Implementation of biochemical reactions in CaiNet. (a) Sketch of an exemplary biochemical reaction comprised of two basic enzyme turnover reactions (upper panel). The two enzymes mutually take each others product and catalyze it to each others substrate. During each time step, molecule numbers and molecule fluxes need to be synchronized to ensure mass conservation (lower panel). (b) Comparison of time trajectories of product levels of the biochemical reaction sketched in (a) simulated with CaiNet using two different synchronization time steps Δ*t* (yellow/purple and green/blue lines) with the numerical solution of an ODE solver (dashed blue/dashed red lines). If the time-step Δ*t* is too large, the transient behavior of CaiNet deviates from an ODE-solver. If the time-step Δ*t* is sufficiently small, the transient behavior of CaiNet approximates the ODE-solver well.

In Equations (8) and (9), the molecule numbers and fluxes that are synchronized at the end of each synchronization time step *Δt* are indicated with a circumflex accent. These parameters stay constant during each synchronization time step. With this assumption, we were able to decouple the differential equations of both bi-molecular elements and solve them separately (Methods). The resulting product numbers are reported as output of the bi-molecular elements to all other network elements after each synchronization step *Δt*. We tested how the duration of the synchronization time step *Δt* influenced the accuracy of the results using a fixed set of reaction rates (Table 3) and *Δt* = 1s or 10s (Figure 4b). Similar to the case of gene elements, if *Δt* was too large, synchronization of reaction products was delayed compared to a global ODE solution and approximation of the transient behavior of the global ODE solution was poor. However, the steady state level was still reproduced correctly independent of the size of the time step, since the correct system of differential equations was used to calculate the outputs of the bi-molecular reaction elements. If *Δt* was sufficiently small, the approximation of CaiNet approached the global solution of the associated system of ODEs using a numerical solver.

**Table 3:** Kinetic parameters of the exemplary biochemical reaction

| Paramter | Values |
| --- | --- |
| Enzyme 1 |  |
| $\lambda$ | 0.001 |
| $\mu$ | 0.1 |
| $V$ | 0.2 |
| $\delta$ | 0.001 |
| Enzyme 2 |  |
| $\lambda$ | 0.001 |
| $\mu$ | 0.1 |
| $V$ | 0.1 |
| $\delta$ | 0.001 |

To ensure a sound consideration of molecule abundance using the flux among elements, We implemented that CaiNet returns a warning once the difference in flux *Δj* between start and end time of the synchronization interval violates

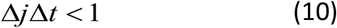

### Inference of GRNs with a neural network approach

We took advantage of the modular approach of CaiNet to facilitate inference of kinetic rates of network elements and the topology of a GRN by using a gradient method originally designed to train recurrent neural networks. The analogy between GRNs and neural networks (NNs) has been pointed out before, by discussing genes as information processing units^43–46^. Accordingly, we directly identified each network element and its divers physiological steady-state input and output parameters with a unique perceptron. Thus, we could directly interpret the GRN laid out in CaiNet as a NN. This identification enabled us to apply a training method designed for recurrent NNs to the GRN, and the process of training corresponded to the process of inference of network parameters. As experimental ground truth, several known steady state measurements of gene product levels including different input conditions of the unknown network or knockout experiments of one or several genes may be taken. During the inference process, we minimized the difference between the current network behavior and the experimental ground truth by optimizing steady-state parameters of the GRN. In particular, we used a back-propagation algorithm introduced for recurrent NNs, which does not perform a global gradient descent step for all nodes in the network, but rather performs a local gradient decent and propagates remaining errors to other nodes^54^ (Methods). The modular layout of the GRN in CaiNet resulting from our central assumption facilitated calculating this gradient.

For an initial proof-of-concept, we applied the process of inference to a network of three genes, for which we assigned a set of reaction rates (Table 4) and activating or repressing connections between the genes (Figure 5a). We then used CaiNet to simulate the ground truth behavior of the GRN for two different inputs. The resulting expression levels served as measurements for the proof-of-concept inference. For simplicity, we only inferred the equilibrium constants of transcription factor binding, i.e. of on- and off-switching of the genes, while leaving production and degradation rates fixed. We assigned false starting values for the equilibrium constant of each gene (Figure 5a), simulated the resulting network behavior and calculated the difference to the two measurements of the ground truth network. Next, we iteratively applied the adapted gradient method until the difference between simulated and ground-truth values dropped below a certain threshold (Methods). During each iteration, the parameters in the guessed network were changed such that the difference between simulated and ground-truth values became smaller. These changes of parameters propagate through the network. Due to feedback loops in the network structure, such a change may cause worse performance of the guessed network. Therefore, the distance between ground truth and guessed network may oscillate before reaching a stable value (Figure 5b–d). The resulting values for the equilibrium constants and thus the performance of the network were well regained during the inference process (Figure 5a).

**Table 4:** Kinetic parameters of the exemplary GRN used to demonstrate the network inference

| Paramter | Scenario: Slow Switching | Equilibrium constant |
| --- | --- | --- |
| Gene 1 |  |  |
| $\lambda_A$ | 0.01 | 0.1 |
| $\mu_A$ | 0.1 | |
| $\lambda_R$ | 0.01 | 0.1 |
| $\mu_R$ | 0.1 | |
| $V$ | 0.1 | |
| $\delta$ | 0.001 | |
| Gene 2 |  |  |
| $\lambda_A$ | 0.01 | 0.1 |
| $\mu_A$ | 0.1 | |
| $V$ | 0.1 | |
| $\delta$ | 0.001 | |

**Figure 5:**
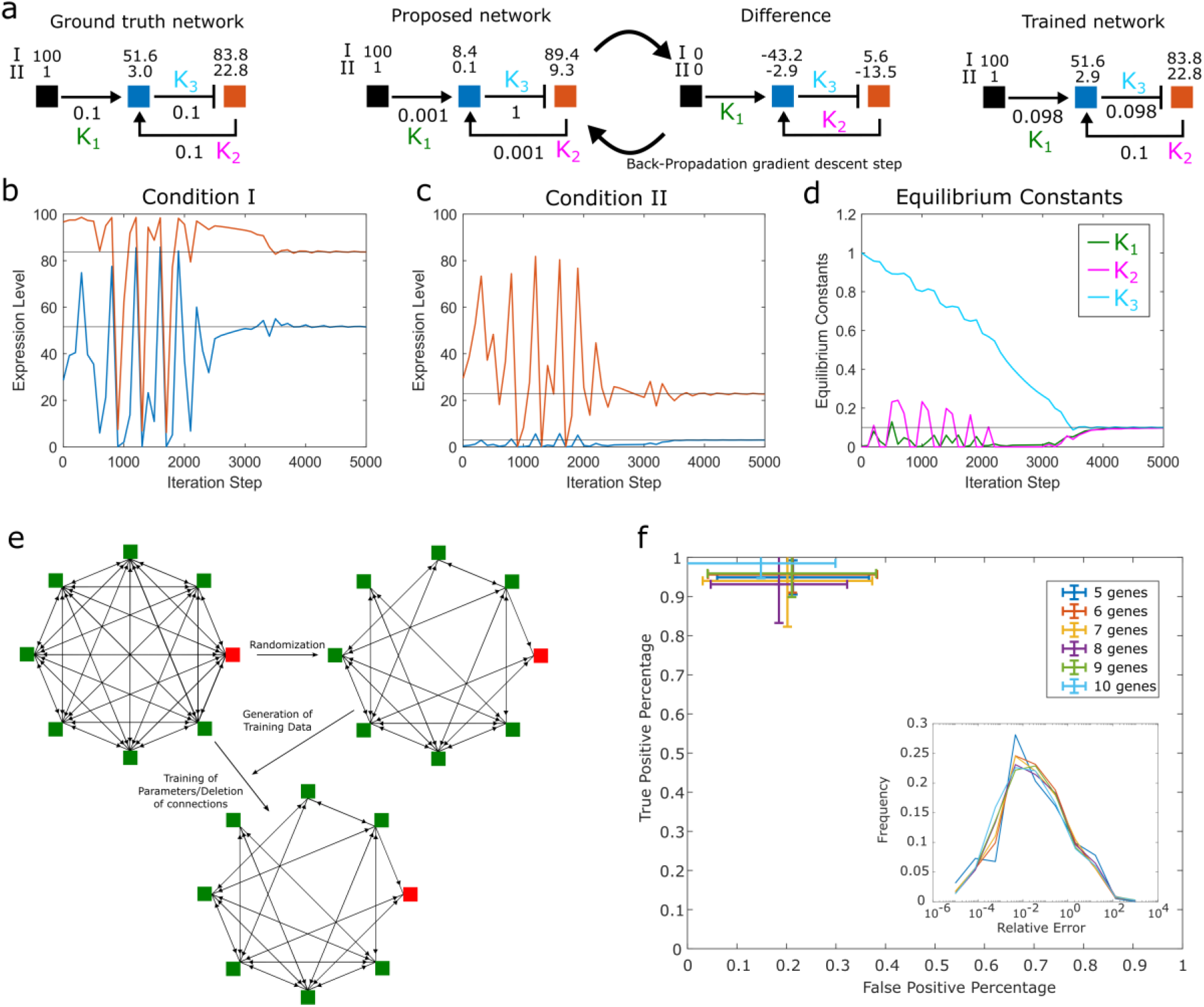
Inference of GRNs with CaiNet using a recurrent network training approach. (a) Demonstration of the inference procedure using a GRN comprising an input element and 2 genes. *Left panel:* Sketch of the ground truth network. The equilibrium constants of transcription factor binding and gene product levels resulting from two different input levels (I and II) are depicted. *Middle panel:* Sketch of the first training cycle. Based on the difference between a guessed network and the ground truth network and the gradient of the gene response functions, CaiNet iteratively adapts the parameters of the network until the expression levels of the guessed network match the ground truth values. *Right panel:* Sketch of the trained network. Trajectories of the gene product levels of gene 1 (blue) and gene 2 (red) for input level I (b) and input level II (c). (d) Trajectories of equilibrium constants of transcription factor binding. (e) Sketch of the procedure to evaluate the inference approach. A ground truth network is generated by randomly omitting gene connections of a fully connected network and assigning equilibrium constants of transcription factors. Gene product levels simulated for the ground truth network are used to train a fully connected network. (f) Percentage of true positive gene connections (agreement between trained and ground truth network) versus percentage of false positive gene connections (exist in trained but not in ground truth network) for GRNs of 5 to 10 genes. Error bars denote s.d. of the training results of 12 different randomly generated networks. *Inset:* Histogram of relative errors (normalized difference in equilibrium constants between trained and ground truth network).

Next, we applied our inference approach to multiple different networks with *N*=5 to 10 genes. To generate these networks, we set up fully connected networks with *N* genes, i.e. that have 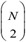 connections (Figure 5e). To obtain a specific ground truth network, we randomly modified the fully connected network. For each connection, we determined whether it was deleted by comparing a random number with a deletion probability *pdelete*, such that on average 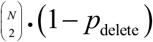 connections remained. For each equilibrium constant of transcription factor binding, i.e. of on- and off-switching of a gene, we drew a random value out of a probability distribution spanning over two orders of magnitude between 1e-3 and 1e-2. As a result of these two steps, we obtained a series of networks with *N* genes and randomly chosen topology and gene switching kinetics, which served as ground truth networks.

To generate measurements for the inference process, we used CaiNet to simulate the expression levels of each gene of a network under different conditions. Here, we knocked out each gene in a network one at a time and simulated the resulting expression levels of the other genes. We repeated this process for each network in the series of networks with *N* genes. We then trained the fully connected, non-randomized, starting network with the gradient descent method on a measurement dataset of a specific network (Methods). During the inference process, CaiNet did not explicitly delete connections. Rather, the equilibrium constants of unneeded or contradictory connections approached zero. We interpreted a connection as deleted, if its equilibrium constant was smaller than 1e-4. With equilibrium constants smaller or equal to this value, the affinity of the corresponding transcription factor is too weak to regulate the expression level of the respective gene. We repeated this process of inference for each network in the series of networks with *N* genes.

To quantify the performance of the inference approach, we counted the number of true positive and false positive connections of a trained network, as compared to the ground truth network topology (Figure 5c). For networks of *N*=5 to 10 genes, the false positive fraction was on average ≈20%, and the true positive fraction was ≈90%. We further quantified the inference of the equilibrium constants by calculating the difference between ground truth and inferred value and normalization this with the ground truth value (Figure 5c, Inset). On average, the relative error of equilibrium constants was ≈1%.

## Disucssion

CaiNet is designed to assist users in setting up and simulating complex GRNs at molecular detail. Simplifying the process of setting up networks, we designed gene elements to assume a two-state behaviour with off and on state of the gene and with a variable number of successive production and processing rates and a degradation rate of the gene product. After assigning the promoter structure and number of successive gene synthesis rates, CaiNet assigns the corresponding local analytical solutions or stochastic models to the gene elements without further input by the user. This is possible since we treat each gene element isolated from other network elements between two synchronization time steps.

Our modular algorithm enables adding more and more realistic kinetic behaviour and noise in molecule abundance to initial deterministic solutions, since the local analytical treatment of the two-state model and gene product synthesis of a gene element can be readily replaced by a stochastic treatment of gene on/off switching and product synthesis for each gene. Thus, it is possible to quickly assess the effect of stochasticity on the behaviour of a network. Another great advantage of the modular approach of CaiNet is that new GRNs that follow the implemented regulatory structure can be set up and simulated very quickly. Another advantage is that simulations of the modular stochastic-deterministic elements can be fast compared to the Gillespie method.

The precalculated analytical solutions applied in CaiNet set a limit for the generality of modelling biological networks. For now, we implemented modelling of genes only with two states and a few promoter structures. In principle, new network elements with further promoter structures or more complex multistate models can be implemented in CaiNet as specialized effective two-state models. It has been found that such effective models are suitable to well describe the histogram of mRNA levels of a cell population^55^. We further assumed a direct relationship between transcription factor binding and activation of a gene. Epigenetic alterations to the gene locus or further steps such as recruitment of the transcription machinery are not modelled explicitly.

While CaiNet is optimized for setting up GRNs and simulating stochastic processes in gene product synthesis, biochemical reactions can also be implemented to account for molecular signalling pathways and reaction cascades that are oftentimes connected to GRNs. For these pathways, we did not implement stochastic fluctuations in the expression levels, since biochemical reactions typically are fast compared to the kinetics of gene on/off switching. For biochemical reactions, CaiNet’s elements are limited to dimerization reactions and enzyme-mediated transformations of biomolecules. These basic bimolecular reactions can be combined to more complex biochemical reactions. However, biochemical reactions with more complex response functions or hill coefficients larger than two are not implemented and require calculating new CaiNet elements.

CaiNet is able to infer GRNs from steady state expression levels. The inferred parameters are therefore limited to equilibrium constants. Since it is challenging to define a gold-standard for inference methods^42,56^, we refrained from comparing CaiNet with other inference methods, such as previously attempted in the DREAM challenge. Instead, we used a comprehensive randomized approach to evaluate the performance of CaiNet. Our simulations indicate that if each gene in a network is knocked out individually, the resulting gene product levels provide sufficient data to infer the topology and parameters of the network. A limit for the size of the inferred network is given by the computation time of the recurrent network algorithm and by the amount of known features of the GRN. Importantly, the inferred equilibrium constants directly correspond to physiological parameters of the GRN and therefore directly allow for subsequent forward simulation of the inferred GRN.

## Conclusion

We provide a user-friendly framework, CaiNet, to simulate GRNs at molecular detail and to infer the topology and steady-state parameters of GRNs. In CaiNet, biochemical reactions and genes with various regulatory logics can be combined to complex GRNs, for which the temporal progression of gene product levels is simulated. Conversely, experimental steady-state measurements of expression levels can be used to infer the underlying GRN. The combination of a forward simulation tool and inverse inference tool in the single package CaiNet may facilitate the process of evaluating biologically relevant GRNs and interpreting associated measurement results.

## Methods

### Pseudo code of network simulations by CaiNet

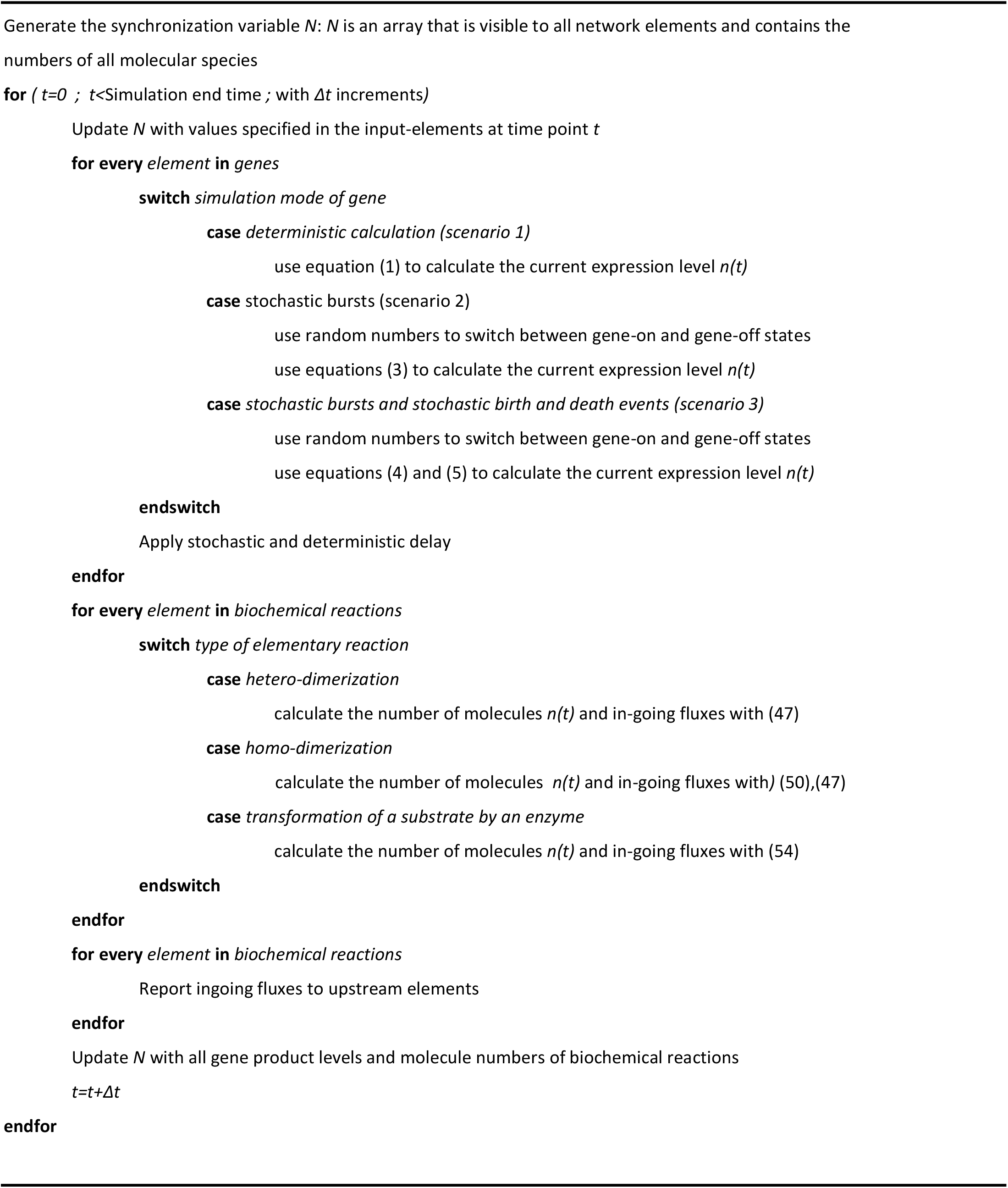

### Effective rates given by the promoter structure

#### Activation by a single transcription factor

For the most simplified case of activation of a gene we assume that a promoter is activated once a transcription factor (TF) binds. This means that the on-rate is given by the arrival rate *λ_eff_* of the TF at the promoter. Once a TF has arrived at the Promoter the gene is ‘on’ for the time 1/*μ_eff_*. This time may vary depending on the TF and the promoter of the gene. If not otherwise stated we assume that the binding time of the TF, 1/*μ*, corresponds to this on-time.

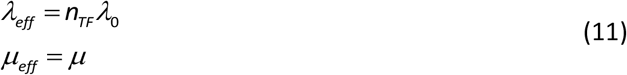

where *n_TF_* is the number of transcription factors and *λ*_0_ is the arrival rate of a single transcription factor.

#### Promoter with AND-logic

The AND-logic refers to a promoter that is only activated if TF_1_ and TF_2_ up to TF_N_ are bound (Figure 2d). Once a single TF leaves, the activation criterion is immediately violated and the promoter is off. Therefore, the off-rate *μ_eff_* of the promoter is the sum over all off-rates *μ_TFi_* of the TFs.

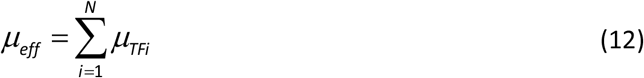

Combinatorically, we can also write down the probability of the promoter to be on as the product of the probability of all TFs to be bound

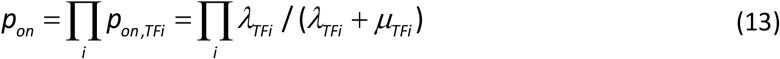

where *λ_TFi_* = *λ_TFi,0_ n_TFi_* is the arrival rate of a TF at the promoter. From *p_on,eff_* and *μ_eff_*, we can calculate the on-rate of the promoter

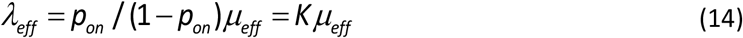

#### Promoter with OR logic

When a promoter is active if TF_1_ or TF_2_ up to TF_N_ or a combination of all is bound, we refer to this promoter as OR-logic (Figure 2d). We start the calculation of effective rates with the on-rates of individual TFs, *λ_TFi_*. Since the arrival of any TF is enough to activate the promoter, the on-rate is

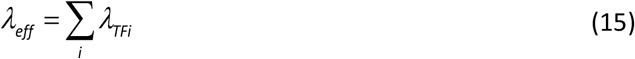

The probability to be on can be calculated combinatorically with *pon,TFi* implicitly defined in (13)

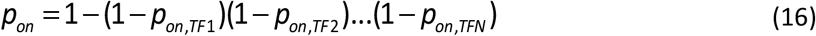

With (14) we find the effective off rate of the promoter

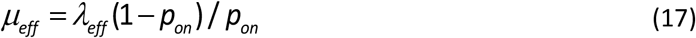

#### Competitive Repression

We now enhance all promoters developed above by an additional feature, which is the blocking of the promoter by a repressor. We assume that the promoter cannot be activated once a repressor is bound. Once a gene is activated however, the repressor does not abort expression. Therefore, the repressor directly affects the effective on-rate of an arbitrary promoter while the off-rate remains unchanged

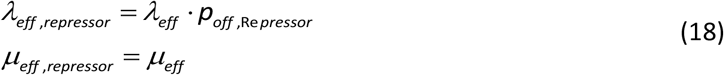

To proof this, we write down the differential equations describing the change in the blocked and free promoter populations. We denote blocked promoters with *b*, free but inactive promoters with *f* and active promoters with *a*

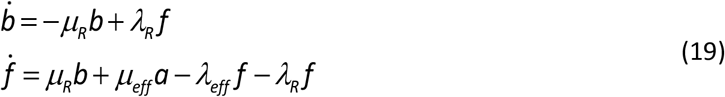

Assuming that the changes in the repressor-population are small, we find

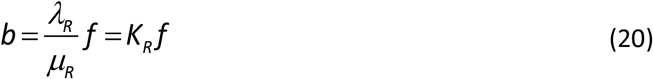

If we now calculate the sum of blocked and unblocked promoters, we obtain

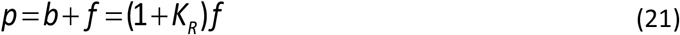

We add up equations (19), plug the result in (21) and obtain

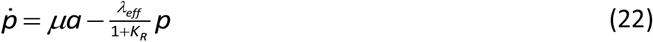

From this result we conclude for the new effective rate modified by the repressor

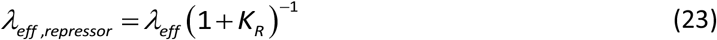

This is equivalent to Equation (18) and the proof is complete.

#### Gene expression including delays

In a simplified approach, we considered one rate-limiting step with rate constant *ν* for synthesis of the gene product. This assumption neglects the fact that certain processes in gene product synthesis may introduce a delay between initiation of product synthesis and availability of the synthesized product, e.g. elongation of RNA, termination of transcription, initiation of translation, elongation of the protein and termination of translation. In addition, the mRNA needs time to be transported to ribosomes and might be subject to posttranslational modifications. To account for such processes in gene product synthesis, we included additional rate-limiting steps corresponding to Poisson processes with rates *β*_1_…*β*_N_.

To determine the resulting function *n(t),* we solve the general system of ODEs for an arbitrary number of delay processes. Using the switching function *a(t)* of a single gene, we arrive at the system of differential equations

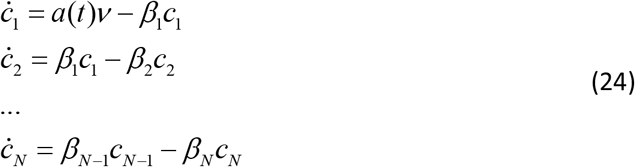

where *c_N_* represents the output expression level of the gene. Our first step to solve the system for *c_N_(t)* is to calculate the Laplace transform of all equations in (24). The result for the first and the i-th equation is

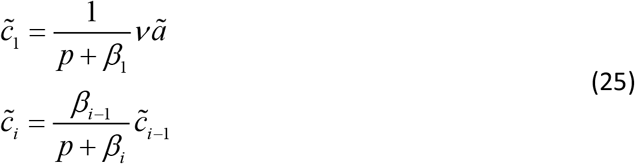

where *p* the transform variable and 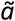 is the Laplace transform of *a*. With these results we can easily find an equation for the Laplace-transform of the final product *c_N_*

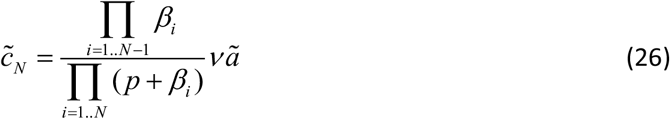

Before we calculate the inverse Laplace transform to obtain *c_N_(t)* we simplify the denominators in (26) by partial fraction composition with the coefficients *α_i_*

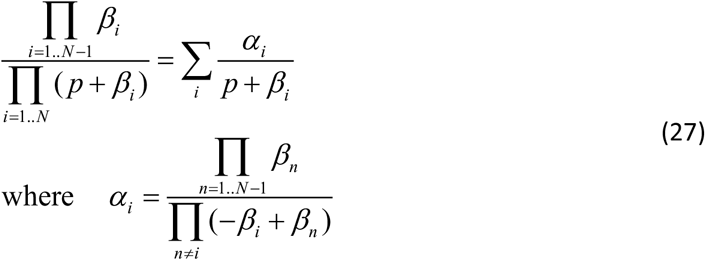

For few delays this simplification can most easily be verified by plugging in *α_i_* and rewriting the right hand side of (27). For a high number of delays the result can be derived by the well-known technique of partial fraction decomposition. We plug this result in (26) and obtain

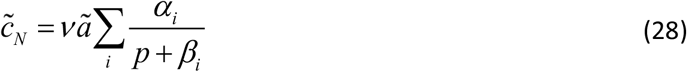

We next use the multiplication theorem for the Laplace transform and find

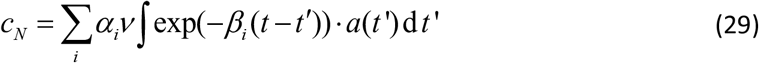

#### Training of recurrent regulatory networks

We assume that steady state expression levels for a subset of species of a GRN are given. Our goal is to infer parameters characterizing the steady state behaviour of the network from this information. Importantly, since we work with steady state information, our approach is limited to the inference of equilibrium constants rather than kinetic rate constants. To infer these constants, we apply a gradient method particularly design for recurrent network topologies^54^. Once these equilibrium parameters are inferred, the temporal evolution of the network can still be simulated.

We start by writing down the minimization problem for the cost function *W(ϕ)* of the difference between the current values ***X*** and the target expression levels ***T*** of the network. The cost function shall be minimized by varying the parameters of the GRN, *ϕ* = {*λ*, *μ*, *ν*, *δ*}.

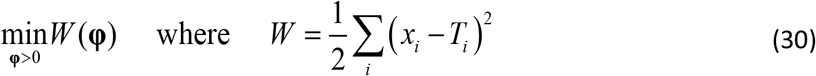

To obtain the gradient for a parameter *ϕ_j_* ∈ ***ϕ*** we calculate the derivative of the cost function with respect to *ϕ_j_* and obtain

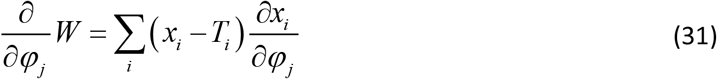

Using the delta rule introduced by Pineda et al.^54^, we find a step size for the parameter *ϕ_j_*

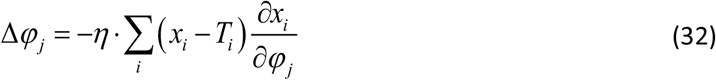

where we use the empirical parameter *η* to scale the step size. In the above equation, the derivatives of *x_i_* with respect to *ϕ_j_* are unknown. To derive a formula for *Δϕ_j_*, we in the following formally work on a generic system of ODEs corresponding to a GRN.

The time evolution of all expression levels ***X*** = *{x_1_, …, x_i_}*, is given by the set of differential equations 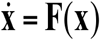. To obtain the missing derivatives we take the total derivative with respect to *ϕ_j_* of the steady state case *0* = **F(*X*)**. We obtain

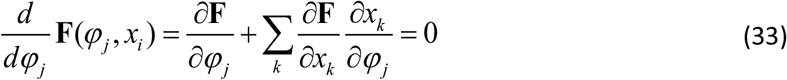

In this equation we recognize the partial derivative of *x_k_* with respect to *ϕ_j_*. Importantly, the system of equations is linear with respect to this derivative. We introduce the abbreviation

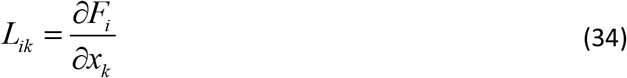

With this abbreviation, we can rewrite (33) as a linear system of equations for the derivatives of *x_k_* with respect to *ϕ_j_*.

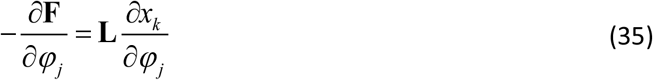

By introducing a new variable *z*, we can make the step of solving this system of ODEs independent from the parameter *ϕ_j_*, such that we only need to solve the system once in order to obtain the gradients for all parameters in ***ϕ***. This variable *z* is characterized by the equation

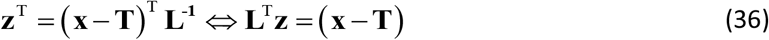

We plug this equation in (32) and identify *z* by introducing the identity matrix. We obtain

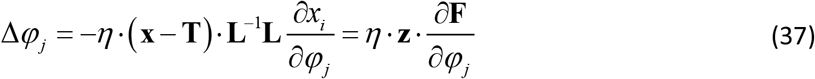

To calculate this gradient using a network set up in CaiNet, we need to calculate the derivatives of the differential Equations 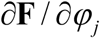 and the derivatives 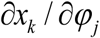. Since the parameter *φ_j_* and the species *x_i_* only occur in a single network element, each network element can return these derivatives independently of other elements in the network.

For a promoter with or-logic, the differential equation is

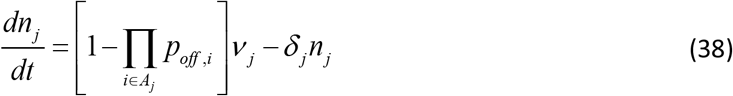

Where *A_j_* is the set of all transcription factors that activate the element *j*. We now take the derivative of the steady state equation with respect to *λ*_k_ and obtain

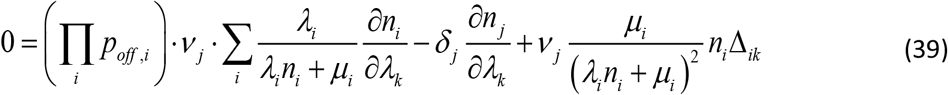

where Δ*_ik_* is the Kronecker delta. From this equation we can identify the variables that the network element returns to calculate a gradient decent step:

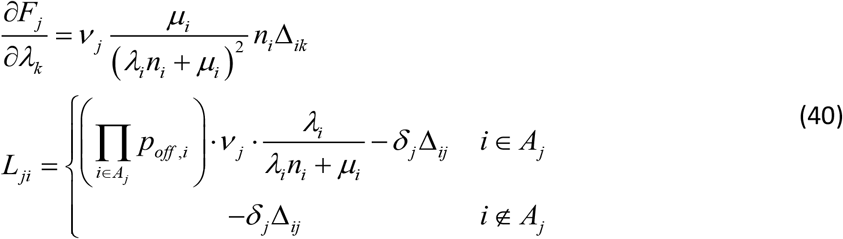

For a complete gradient descent scheme, we first obtain the steady state expression levels by simulating the network with CaiNet until all elements have reached a steady state. Next we use equation (37) to calculate the change for all parameters in ***ϕ*** based on the derivatives above and the difference between ground truth expression levels and simulated expression levels. Using the new parameters, we again simulate the steady state expression levels. We proceed in this manner until an abortion criterium is met. This abortion is *W* < *b*, i.e. that the sum of all differences between ground truth and expression levels of the trained network are smaller than an upper bound *b*.

#### Chemical Reactions

In the following we derive analytical solutions for the elementary reactions ‘homo-dimerization’, ‘hetero-dimerization’ and ‘transformation of a species by an enzyme’. We then use these analytical solutions to couple multiple elementary reactions to obtain more complex biochemical reaction networks.

#### Hetero-dimerization

We start with the hetero-dimerization of the two species *n*_1_ and *n*_2_ that forms the species *f*. For a closed system of these three species the change in *f* over time is

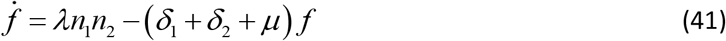

where *λ* is the association rate of the two dimerizing species and *μ* is the dissociation rate of the dimer. The rates *δ*_1_ and *δ*_2_ correspond to the degradation of the monomers *n*_1_ and *n*_2_ respectively. The law of mass conservation yields equations for the species *n*_1_, *n*_2_

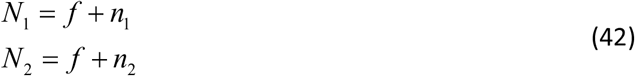

Plugging in the law of conservation we obtain

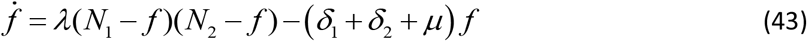

To account for coupling to other elementary systems, we introduce the flow *j_in_* that represents flux of the species *f* from other elements into the element at hand. The new differential equation for *f* is

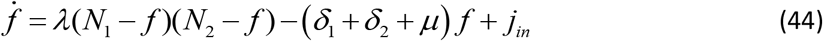

Next, we solve this equation for *f* (*t*) for an initial value of *f*_0_. We start by calculating the fixed points 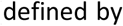 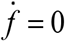 of *f*. The corresponding quadratic equation yields the fixed points *f*_1_, *f*_2_

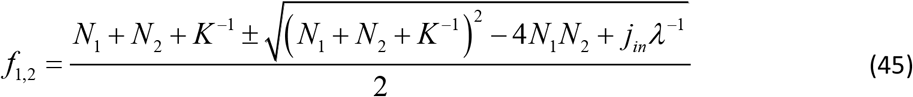

where *K* = *λ*· (*μ*+*δ*_1_+*δ*_2_)^−1^. Using these fixed points, we can rewrite equation (44) as

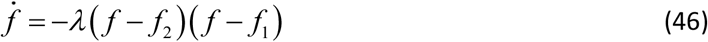

Integration of this ODE yields

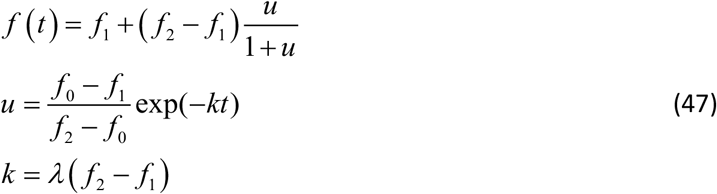

Since the reaction consumes the species species *n*_1_, *n*_2_, we need to calculate the flux of the respective elements into the species *f*. This flux is calculated by

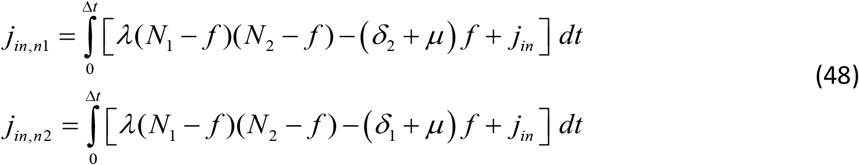

During each synchronization time-step, these fluxes are reported to the network element representing the corresponding species.

#### Homo-Dimerization

The differential equations for a homo-dimerization are

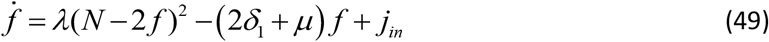

where *λ* is the association rate of the two dimerizing monomers and *μ* is the dissociation rate of the dimer. The rate *δ*_1_ corresponds to the degradation of the monomers *N*. The shape of this equation is similar to the heterodimer case. Thus, by setting

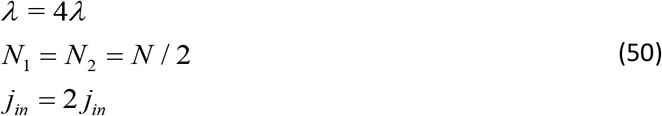

we can reuse our solution (47) to calculate the number of dimers and (48) to calculate the flux out of the monomer into the homodimer. During each synchronization time-step, this flux is reported to the monomer element.

#### Enzyme kinetics

In case of a dimerization with an enzyme, the substrate can be subject to a reaction catalysed by the enzyme. Here, the species *m* is produced.

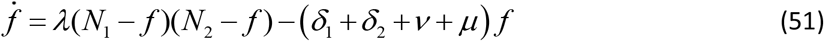

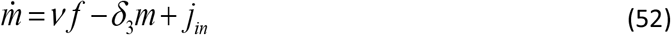

where *λ* is the association rate of substrate to enzyme and *μ* is the dissociation rate of the substrate-enzyme complex. The rates *δ*_1_, *δ*_2_ and *δ*_3_ correspond to the degradation of the substrate *n*_1_, degradation of the enzyme *n*_2_ and degradation of the product respectively. The rate *ν* is the transformation rate of substrate to product. The shape of this equation is similar to the heterodimer case. Thus, by setting

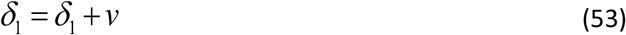

we can reuse our solution (47) to calculate the number of enzyme-substrate-complexes and (48) to calculate the flux out of substrate and enzyme into the enzyme-substrate-complex. To obtain the amount of product, we integrate (52) and obtain

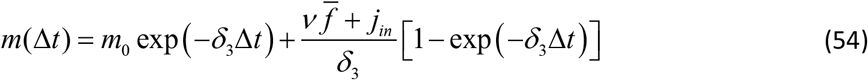

where

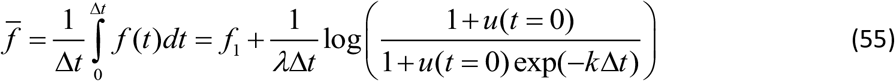

We arrived at this result by reusing our solution (47). During each synchronization time-step, *m(Δt)* is reported to all other elements in the network.

### Simulation of the negative feedback Cascade

#### Gillespie Simulation

All genes in the cascade have three states, since the promoter structure is not modelled with an effective two-state model as in the case of the CaiNet simulation. The promoter is empty in the second state. By binding of a transcription repressor the promoter enters the first state. This state can only be exited upon unbinding of the transcription repressor. The third state is entered upon binding of a transcription activator

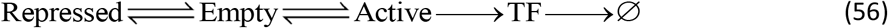

This state model leads to the following system of stochastic reactions. We consider the n-th link of the chain *P_n_* then is transcriptionally repressed by the gene product of the (n-1)-th link. The n-th link of the chain produces the transcription repressor *R_n_*

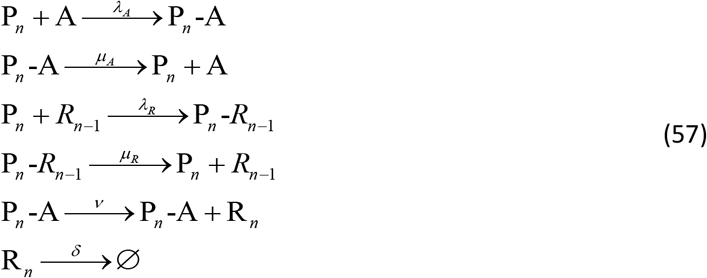

We implemented this set of reactions for *n* = 1,2,3,4.

#### System of ODEs

For the system of ODEs we do not explicitly simulate the association and dissociation of transcription factors to the promoters. Rather, we give the on-probability of the gene. The state model (56) corresponds to a competitive repression model of a promoter and we can give the on-probability of the n-th link of the chain using (18)

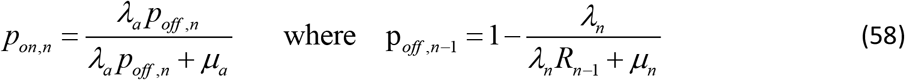

With this on-probability we can give the ODE for the gene product

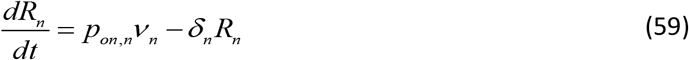

We implemented this set of reactions for *n* = 1,2,3,4. For solving the system of differential equations we used the ode45 solver of Matlab 2019a.

## Acknowledgements

Not applicable.

## Author’s contributions

J.H. and J.C.M.G. designed the study; J.H. performed calculations and programmed CaiNet; J.H. and J.C.M.G. wrote the manuscript.

## Availability of data and materials

The CaiNet software is freely available at https://gitlab.com/GebhardtLab/CaiNet

## Declarations

### Ethics approval and consent to participate

Not applicable.

### Consent for publication

Not applicable.

